# Dysregulated tissue-resident lymphocytes drive senile emphysema by impairing alveolar regeneration

**DOI:** 10.64898/2026.04.22.720146

**Authors:** Yanjing Su, Xinguo Yang, Ziyi Ren, Yuxuan Guan, Xuelu Zhou, Shuilian Chi, Yanjun Huang, Tao Yan, Jialong Liang, Fei Gao, Dijun Chen, Jingyu Chen, Zimu Deng, Chaoqun Wang

## Abstract

Aging is often associated with progressive tissue degeneration and chronic inflammation, yet the role of immune cells in mediating structural and functional decline in organs remains poorly defined. Here, we investigated immune-tissue interactions in the aged lung and identified emphysematous remodeling characterized by alveolar loss. Notably, aged lungs exhibited a marked expansion of tissue-resident lymphocytes (TRLs) with senescent features, accompanied by a significant reduction in alveolar stem/progenitor cell (AT2) abundance. In vivo adoptive T cell transfer and 3D immune-stem cell organoid assays revealed that these expanded TRLs suppressed AT2 growth via secretion of oncostatin M and interferon gamma. In vivo blockade of IL-7 receptor (IL-7R) reduced TRL accumulation in the lungs and ameliorated age-related emphysematous changes, including restoration of alveolar density. Our findings identify TRLs as key drivers of alveolar degeneration in aging and propose IL-7R inhibition as a therapeutic strategy to mitigate pulmonary decline.

**Teaser:** Blocking IL-7R clears harmful lymphocytes and helps rebuild the damaged air sacs of the aging lung.

## INTRODUCTION

Aging drives progressive functional decline across organs, including the lung, where pulmonary performance begins to diminish as early as the third decade of life (*1*). Lung aging is accompanied by a range of molecular, cellular, and physiological alterations that compromise tissue repair and regenerative capacity, thereby increasing susceptibility to chronic pulmonary diseases such as emphysema (*2, 3*). These changes are associated with stem cell dysfunction and chronic low-grade inflammation, commonly termed “inflammaging” (*4*).

Emphysema is a progressive and destructive lung disease that represents one of the most common manifestations of pulmonary aging (*5*). As a major subtype of chronic obstructive pulmonary disease (COPD), it is characterized by irreversible loss of alveolar structures and chronic inflammation, resulting in impaired gas exchange and progressive decline in lung function. Normal physiological aging is associated with enlarged alveolar spaces and a gradual decline in lung function, a condition known as “senile emphysema” (*2*). Despite its high prevalence, no targeted therapies for emphysema currently exist, and the clinical and economic burden of this disease remains substantial (*6*). Thus, understanding the mechanisms linking aging to the loss of lung function is crucial for developing targeted therapeutic strategies for emphysema.

Alveolar epithelial type 2 cells (AT2s) maintain alveolar integrity by producing surfactant and serving as alveolar stem/progenitor cells for epithelial renewal (*7*). With aging, AT2s progressively lose their regenerative capacity (*8*), but the mechanisms driving this decline remain poorly understood. Tissue-resident immune cells, which inhabit organs such as the lung, play intricate roles in regulating stem/progenitor cell function and maintaining tissue homeostasis (*9, 10*). Aging disrupts their homeostasis, leading to immunosenescence and acquisition of a proinflammatory senescence-associated secretory phenotype (SASP), which sustains chronic inflammation and impairs repair (*3, 11*). While this aged immune state contributes broadly to age-related pathologies (*12, 13*), its specific role in age-related emphysema remains unclear.

Here, we investigate how tissue-resident lymphocytes (TRLs) affect AT2s and contribute to age-related emphysematous changes. Our study demonstrates that the expansion of TRLs impairs AT2 regenerative capacity and drives emphysema in the aged lung.

## RESULTS

### Aged lungs exhibit emphysematous changes and AT2 dysfunction

Naturally aged lungs were characterized by complex, multiscale alterations in tissue architecture, accompanied by changes in cellular and molecular composition (*8*). Here, we examined age-associated morphological changes in alveoli of young and aged human lungs (Table S1).

Histological analysis of aged human lungs revealed marked enlargement of alveolar spaces accompanied by a reduction in AT2 cell abundance, compared with young lungs (Fig. 1, A-C). Consistent with observations in human lungs, analysis of mouse lungs revealed a progressive increase in airspace size and mean linear intercept (MLI) with age, along with reduced alveolar density (Fig. 1, D-G). Immunofluorescent analysis further demonstrated a significant reduction in AT2 cell number in aged lungs (Fig. 1, H and I). Together, these data indicate that hallmarks of age-associated emphysema, characterized by alveolar loss and AT2 depletion, are conserved in both humans and mice.

**Fig. 1.**
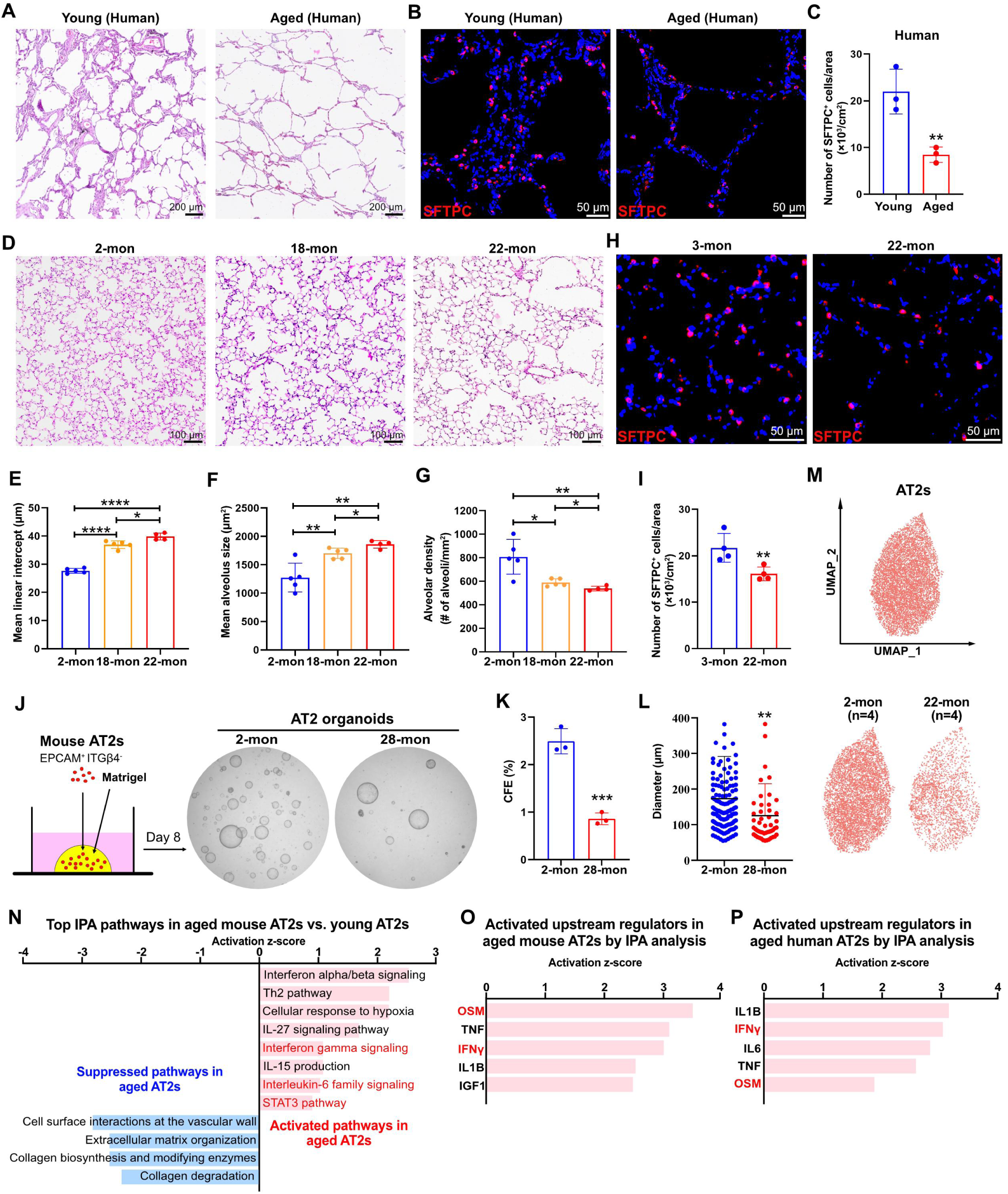
Emphysematous remodeling and impaired stem cell regeneration in aged lungs. (**A**) H&E images of lungs from young (age = 42-45 years) and aged (age = 55-70 years) humans. (**B**) IF analysis of SFTPC in lungs from young and aged humans. (**C**) Quantification of AT2 cell numbers in lungs from young and aged humans. (**D**) H&E images of lungs from young and aged mice. (**E** to **G**) Quantification of MLI, mean alveolus size, and alveolar density in lungs from young and aged mice. (**H**) IF analysis of SFTPC in lungs from young and aged mice. (**I**) Quantification of AT2 cell numbers in lungs from young and aged mice. (**J**) Representative images of organoids derived from AT2s isolated from young and aged mice (CD31^−^/CD45^−^/PDGFRα^−^/EPCAM^+^/ITGB4^−^). (**K** and **L**) Quantification of colony-forming efficiency (CFE) and organoid size. (**M**) scRNA-seq of AT2s from young and aged mouse lungs. (**N**) Activated and suppressed pathways in aged AT2s relative to young mouse AT2s, analyzed with IPA. (**O**) Predicted upstream regulators in aged versus young mouse AT2s, analyzed with IPA. (**P**) Predicted upstream regulators in aged versus young human AT2s, analyzed with IPA. Each data point represents one human or one mouse (**C**, **E** to **G**, and **I**) of an individual experiment. Data are expressed as Mean ± SD. *p < 0.05; **p < 0.005; ****p < 0.0001.

To investigate the mechanisms underlying AT2 loss with aging, we employed a 3D organoid culture system of AT2s (*9*). AT2s isolated from young (2-month-old) and aged (28-month-old) mice were implanted in Matrigel droplets and cultured to assess their regenerative capacity (Fig. 1J and Fig. S1A). Notably, AT2s isolated from aged mice exhibited a markedly reduced capacity to generate organoids, as evidenced by decreased colony-forming efficiency (CFE) and smaller organoid size (Fig. 1, J-L), consistent with previous findings (*14*). Next, we performed single-cell RNA sequencing (scRNA-seq) to examine age-associated transcriptomic changes in AT2s (Fig. 1M and Fig. S1B). Differentially expressed gene (DEG) analysis of AT2s from young versus aged lungs (Table S2), coupled with ingenuity pathway analysis (IPA), revealed activation of immune-related pathways in aged AT2s, including interferon gamma (IFNγ) signaling, Th2 pathway, interleukin-6 family signaling, and STAT3 pathway, most of which were linked to T cell activation (Fig. 1N). These findings suggest that AT2s in aged lungs are subjected to heightened immunological stress, particularly from T cell-derived signals. To identify potential factors driving AT2 alterations in aged lungs, we performed IPA upstream regulator analysis of AT2 DEGs (aged versus young). Consistent with the pathway analysis, the top predicted upstream regulators in aged AT2s were predominantly immune-related cytokines (Fig. 1O), including oncostatin M (OSM), a cytokine implicated in epithelial remodeling and recognized as a biomarker of COPD (*15, 16*), and IFNγ, which we have previously shown to drive emphysematous changes following viral exacerbation (*9*). To investigate whether human and mouse AT2s undergo similar transcriptional alterations with aging, we reanalyzed previously published scRNA-seq data from aged and young human lungs (Table S3 and S4) (*17*) and compared the results with DEGs identified in mouse AT2s (aged versus young). Hypergeometric probability testing showed significant enrichment of overlap genes in DEGs between aged human and mouse AT2s (Fig. S1C). Furthermore, activated upstream regulators in aged human AT2s were largely immune-related cytokines, showing strong concordance with the upstream regulators identified in aged mouse AT2s, including OSM and IFNγ (Fig. 1P). Together, these findings indicate that aging reduces regenerative capacity of AT2s and alters their transcriptional profiles, marked by increased immunological stress, particularly driven by T lymphocytes.

### Resident immune landscape in aged lungs marked by expansion of senescent TRLs

To characterize the tissue-resident immune landscape of aged lungs, we performed intravenous (i.v.) anti-CD45 labeling just prior to lung harvest to differentiate circulating (i.v. anti-CD45^+^) from tissue-resident immune cells (i.v. anti-CD45^−^), as previously reported (*9, 18*). Resident immune cells were then isolated by flow cytometry and subjected to scRNA-seq (Fig. S2A).

Following quality control analyses, filtering, and doublet removal, 89,804 tissue-resident immune cells remained for characterization. Unbiased clustering and uniform manifold approximation and projection (UMAP) analysis revealed 11 distinct resident immune populations, including T cells (*Cd3e*^+^), B cells (*Ms4a1*^+^), natural killer cells (NK; *Ncr1*^+^), innate lymphoid group 2 cells (ILC2; *Il1rl1*^+^), alveolar macrophages (aMac; *Chil3*^+^), proliferating alveolar macrophages (proli aMac; *Chil3*^+^ and *Mki67*^+^), interstitial macrophages (iMac; *C1qb*^+^), mature dendritic cells (maDC; *Fscn1*^+^), classical dendritic cell subset 1 (cDC1; *Clec9a*^+^), classical dendritic cell subset 2 (cDC2; *Cd209a*^+^), and neutrophils (Neu; *S100a8*^+^) (Fig. 2A; Fig. S2, B and C). The composition of lung-resident immune cells was dramatically altered in aged lungs, with T cells exhibiting the most pronounced increase, exceeding a twofold increase (Fig. 2B). These TRLs expressed canonical tissue-residency markers (*19*), including *Il7r* (IL-7 receptor), *Cd69*, and *Cd44*, while lacking expression of circulating markers, such as *S1pr1*, *Ccr7*, and *Sell* (CD62L) (Fig. S2D). The accumulation of TRLs in aged lungs was further confirmed by flow cytometry and immunofluorescence analyses (Fig. 2, C-F).

**Fig. 2.**
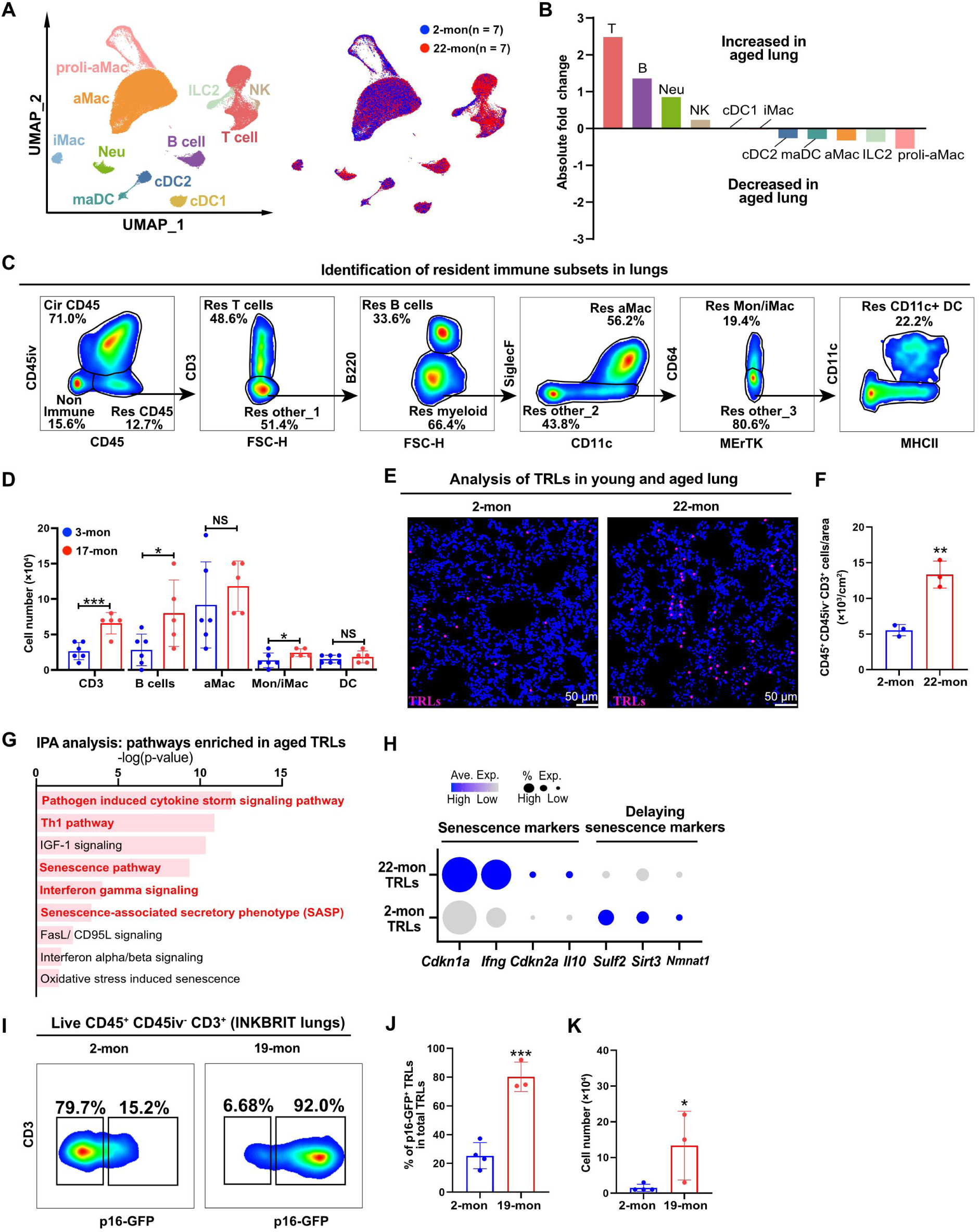
Resident immune landscape featuring expansion of senescent TRLs in aged lungs. (**A**) scRNA-seq of resident immune cells in lungs from young and aged mice. (**B**) Absolute fold change in cell number of the resident immune subsets between young and aged lungs. (**C**) FACS strategy to isolate and quantify resident immune subsets. (**D**) Quantification of the number of resident immune subsets in young and aged lungs by flow cytometry. (**E**) Thick-section confocal imaging of TRLs (CD45^+^/CD45iv^−^/CD3^+^) in the alveoli of young and aged lungs. (**F**) Image quantification of TRLs in young and aged lungs. (**G**) Activated pathways in aged TRLs, relative to young TRLs, analyzed with IPA. (**H**) Plots showing the expression of senescence markers (*Cdkn1a*, *Ifng*, *Cdkn2a*, *Il10*) and markers associated with delayed senescence (*Sulf2*, *Sulf3*, *Nmnat1*) in young and aged TRLs. (**I**) Flow cytometry analysis of *p16*^INK4a+^ TRLs in young and aged lungs. (**J**) Percentage of *p16*^INK4a+^ TRLs in total TRLs, analyzed by flow cytometry. (**K**) Quantification of *p16*^INK4a+^ TRLs in young and aged lungs by flow cytometry. Each data point represents one mouse (**D**, **F**, **J** to **K**) of an individual experiment. Data are expressed as Mean ± SD. *p < 0.05; **p < 0.005; ****p < 0.0001.

DEG analysis comparing TRLs from aged and young lungs (Table S5), combined with IPA analysis, showed that pathways activated in aged TRLs were predominantly associated with antigen responses including pathogen-induced cytokine storm signaling pathway, Th1 pathway, and interferon gamma signaling (Fig. 2G). Notably, the senescence pathway was also activated in aged TRLs (Fig. 2G). Consistently, aged TRLs exhibited elevated expression of canonical senescent markers (*20*), including *Cdkn1a* and *Cdkn2a*, along with SASP-related genes, such as *Ifng* (Fig. 2H). In contrast, genes associated with the attenuation of cellular senescence (*21, 22*), including *Sulf3*, *Sirt3*, and *Nmnat1*, were downregulated in aged TRLs (Fig. 2H). T cell senescence is a recognized hallmark of immune aging (*23*). *p16*^INK4a^, encoded by *Cdkn2a*, is a well-established marker of cellular senescence during aging (*23*). Re-analysis of previously published lung scRNA-seq data of aged INKBRITE (*p16^GFP/+^*) mice, an ultrasensitive reporter of *p16*^INK4a^ expression (*24*), revealed elevated SASP enrichment and upregulation of canonical senescence markers in *p16*^INK4a+^ T cells compared with *p16*^INK4a-^ counterparts (Fig. S2, E and F), establishing this model as a robust reporter of T cell senescence. We next performed flow cytometric analysis of the TRLs in young and aged INKBRITE mice (Fig. 2I). Both the fraction and absolute number of *p16*^INK4a+^ TRLs were significantly increased in aged lungs (Fig. 2, J and K). Collectively, these data suggest that aging drives the accumulation of TRLs in the lung, which adopt a senescent phenotype characterized by elevated *p16*^INK4a^ expression and SASP factor production, potentially contributing to impaired lung regeneration and disrupted tissue homeostasis.

### Age-associated TRLs drive more severe emphysematous changes

Next, we investigated the impact of aged TRLs on lung tissue integrity. To this end, we adoptively transferred splenic T cells from young or aged wild-type mice into young recombination-activating gene 1-deficient (*Rag-1*^−/−^) mice, which lack mature T cells, via retro-orbital intravenous injection (Fig. 3A). To assess T cell colonization in recipient lungs 8 weeks post-transfer, we analyzed the lung tissues by colocalizing CD3 with CD44, a marker of tissue residency (*9, 25*), and quantified CD3^+^/CD44^+^ T cells. Robust engraftment of T cells into the lung parenchyma was observed, with no significant difference in colonization efficiency between cells derived from young and aged donors (Fig. 3, B and C). Notably, histological analysis revealed that adoptive transfer of T cells from either group induced emphysematous changes, characterized by alveolar loss and airspace enlargement, as quantified by increased MLI, enlarged mean alveolus size, and decreased alveolar density (Fig. 3, D-G). Importantly, transfer of aged T cells resulted in more severe alveolar destruction, compared with young T cells (Fig. 3, D-G).

**Fig. 3.**
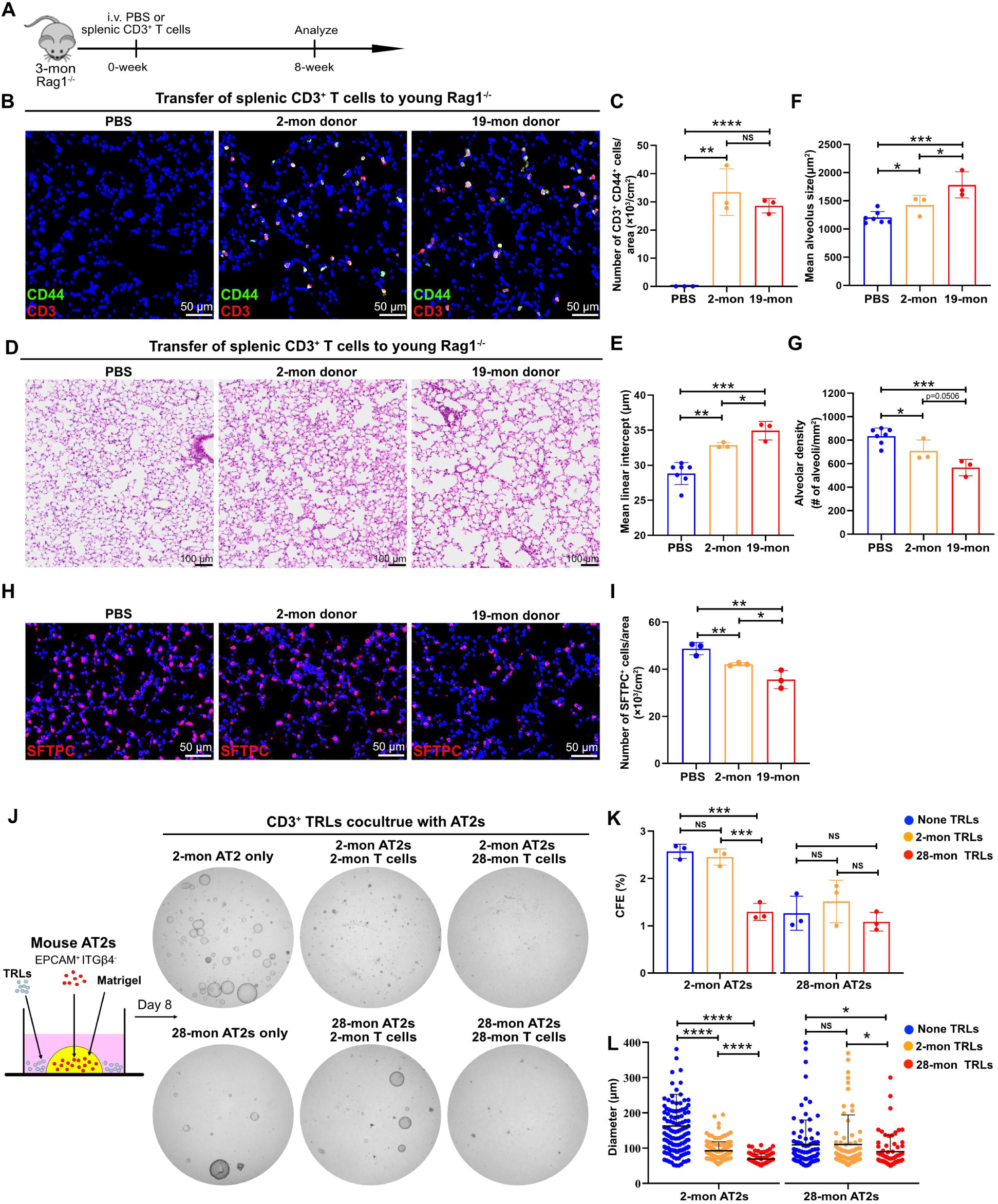
Aged TRLs exacerbate alveolar degeneration and emphysematous pathology. (**A**) Schematic of adoptively transferred splenic T cells from young or aged mice into young *Rag-1*^−/−^ recipients. (**B**) IF analysis of double-positive (CD3^+^/CD44^+^) T cells in *Rag-1*^−/−^ lungs receiving young or aged splenic T cells. (**C**) Quantification of double-positive (CD3^+^/CD44^+^) T cells in recipient lungs. (**D**) H&E images of recipient lungs. (**E** to **G**) Quantification of MLI, mean alveolus size, and alveolar density in recipient lungs. (**H**) IF analysis of SFTPC in recipient lungs. (**I**) Quantification of AT2 cells in recipient lungs. (**J**) Representative images of organoids derived from AT2s co-cultured with TRLs from young or aged mice. (**K** and **L**) Quantification of colony-forming efficiency (CFE) and organoid size. Each data point represents one mouse (**C**, **E** to **G**, and **I**) of an individual experiment. Data are expressed as Mean ± SD. *p < 0.05; **p < 0.005; ****p < 0.0001.

Consistently, a more pronounced depletion of AT2 cells was observed in lungs receiving aged T cells, with a lesser reduction following transfer of young T cells (Fig. 3, H and I).

To determine whether lung TRLs directly suppress AT2 growth, we co-cultured AT2s from young or aged mice with TRLs isolated from young or aged lungs using our immune-stem cell organoid system (*9*). Both young and aged TRLs markedly suppressed the growth of young AT2s, with the inhibitory effect being more pronounced in aged TRLs (Fig. 3, J-L). Aged TRLs also robustly impaired the growth of aged AT2s, whereas young TRLs had little impact, likely reflecting the heightened immune stress already experienced by aged AT2s in vivo (Fig. 3, J-L). Together, these findings demonstrate that TRLs impair AT2 regenerative capacity and drive emphysematous changes, with aged TRLs exerting a stronger inhibitory effect than young TRLs.

### In-depth characterization of TRL subsets in aged lungs

To characterize the age-associated alterations within TRL subsets, we first performed flow cytometry analysis of lung single-cell suspensions prepared from young and aged mice.

Compared with young lungs, aged lungs exhibited a significant accumulation of all TRL subsets, including CD4⁺, CD8^+^, and double-negative TRLs (DNTs: CD3^+^/CD4^−^/CD8^−^) (Fig. 4, A and B). Using *p16^GFP/+^* reporter mice, we further observed that both the percentage and absolute number of *p16*^INK4a*+*^ cells within CD4⁺, CD8⁺, and DNT TRLs were significantly increased in aged lungs (Fig. 4, C-E). These findings indicate a broad activation of cellular senescence programs across all TRL subsets during lung aging.

**Fig. 4.**
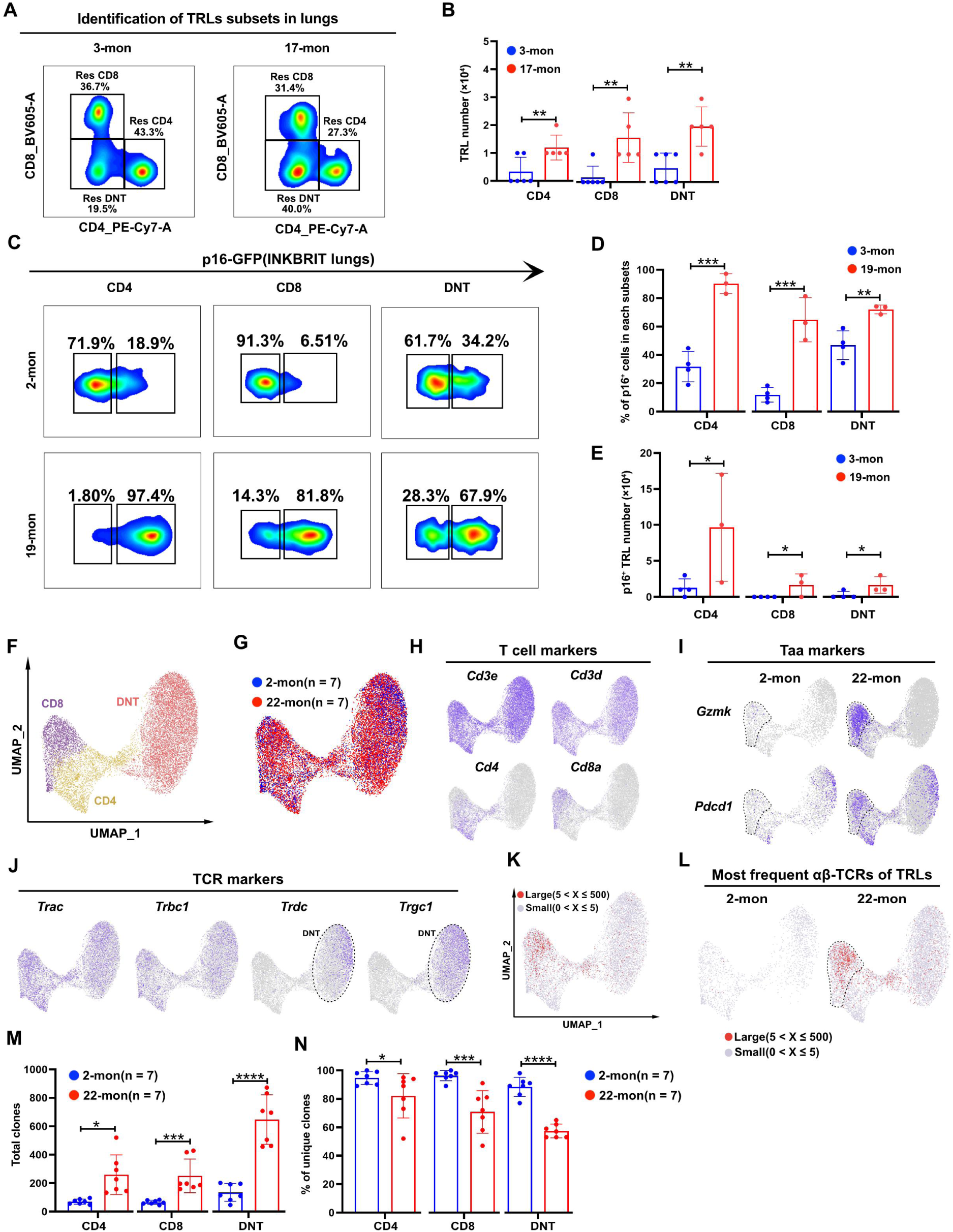
Age-associated alterations in TRL subsets. (**A**) FACS strategy to isolate and quantify TRL subsets in young and aged lungs. (**B**) Quantification of TRL subsets in young and aged lungs by flow cytometry. (**C**) Flow cytometry analysis of *p16*^INK4a+^ cells in TRL subsets from young and aged lungs. (**D**) Percentage of *p16*^INK4a+^ TRL subsets in total TRLs, analyzed by flow cytometry. (**E**) Quantification of *p16*^INK4a+^ TRL subsets in young and aged lungs by flow cytometry. (**F** and **G**) scRNA-seq of TRL subsets from young and aged mouse lungs. (**H**) Plots showing the expression of T cell markers (*Cd3e*, *Cd3d*, *Cd4*, *Cd8a*) in TRL subsets. (**I**) Plots showing the expression of age-associated T cell (Taa) feature markers (*Gzmk*, *Pdcd1*) in young and aged TRL subsets. (**J**) Plots showing the expression of TCR markers (*Trac*, *Trbc1, Trdc, Trgc1*) in TRL subsets. (**K** and **L**) Projection of large (5 < X ≤ 500) and small (0 < X ≤ 5) TCR clones (X, clone size) from young and aged lungs onto TRL clusters. (**M**) Total paired TCRα/β clones in young and aged TRL subsets. (**N**) Percentage of unique clones in young and aged TRL subsets. Each data point represents one mouse (**B**, **D**, **E**, **M**, **N**) of an individual experiment. Data are expressed as Mean ± SD. *p < 0.05; **p < 0.005; ****p < 0.0001.

To further dissect the heterogeneity and TCR repertoire of TRL subsets, we performed scRNA-seq combined with T cell receptor sequencing (TCR-seq). This approach allowed us to comprehensively illustrate the transcriptional profiles and clonotypes of TRLs (Fig. 4, F-H). As shown in Fig. 4I, we found CD8^+^ TRLs of aged lungs markedly upregulated typical markers of age-associated T cells (Taa cells), such as *Gzmk* (granzyme K) and *Pdcd1* (*26*). DNTs, which constitute the largest fraction of TRLs in aged lungs, were primarily identified as γδ T cells (Fig. 4J), marked by the expression of TCRγ (encoded by *Trgc1*) and TCRδ (encoded by *Trdc*) (*27*). In contrast, CD4⁺ and CD8⁺ TRLs exclusively expressed α and β T cell receptors (Fig. 4J). We then analyzed the paired TCRα/β repertoire in TRL populations. Among scRNA-seq-defined αβ T cells, approximately 60% of them were successfully assigned paired TCRα and TCRβ chains (Fig. S3A). TCR clonotype analysis of young TRLs revealed a highly diverse repertoire with low colonality (Fig. 4, K-N), whereas TRLs from aged lungs exhibited a marked reduction in repertoire diversity across all subsets (Fig. 4N). Clonal expansions were observed across all aged TRL subsets, with the most pronounced expansion occurring in CD8^+^ TRLs (Fig. 4L), suggesting dysregulated activation and proliferation of these cells during aging. Together, these data demonstrate a broad expansion of all TRL subsets, including CD4⁺, CD8⁺, and DNT TRLs, in aged lungs. Moreover, aged TRLs exhibit age-associated hallmarks, including upregulation of markers for Taa cells, accelerated cellular senescence, and reduced TCR repertoire diversity.

### TRL-derived IFNγ and OSM inhibit alveolar organoid growth

Having established a functional role of aged TRLs in impairing regenerative capacity of AT2s, we sought to elucidate the mechanisms underlying this effect. We utilized the CellChat package (*28*) to identify enriched ligand-receptor pairs and to predict intercellular communications from TRLs to AT2s. In general, all three TRL subsets exhibited robust communications with AT2s, with DNTs displaying the strongest interaction strength (Fig. 5A). When we looked into individual signaling pathways, we found that CD4^+^ and CD8^+^ TRLs were strongly enriched in IFNγ signaling (Fig. 5B), which has previously been reported to inhibit AT2 growth (*9*). In contrast, DNTs were predominantly enriched for OSM signaling (Fig. 5B). In line with CellChat findings, both OSM and IFNG were identified as the top upstream regulators in aged AT2s (Fig. 1O).

**Fig. 5.**
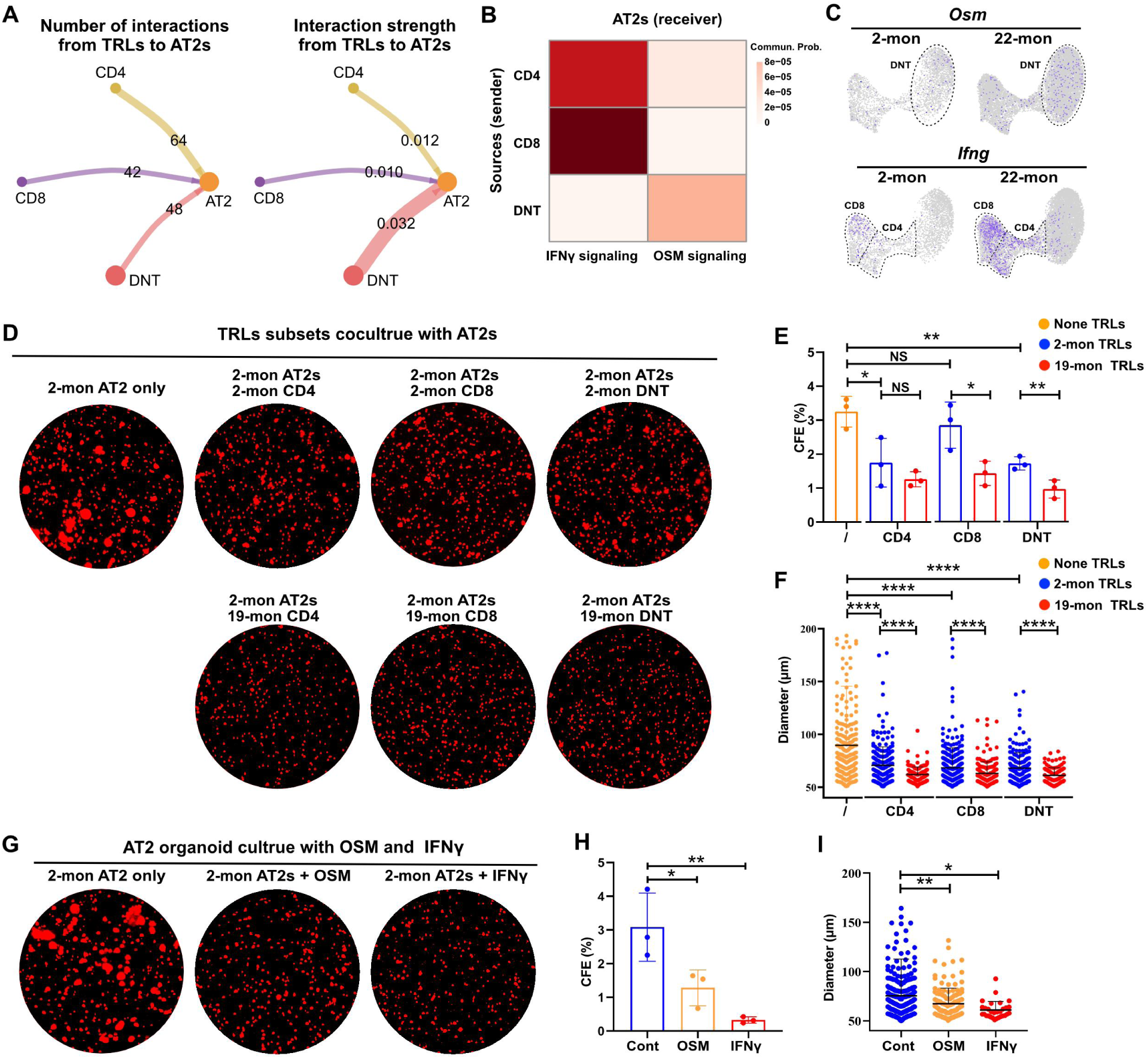
TRL-derived IFNγ and OSM inhibit the regenerative capacity of alveolar organoids. (**A**) CellChat analysis of TRL subset interactions with AT2s. (**B**) Heatmap showing the interactions of IFNγ and OSM signaling pathways from TRL subsets to AT2s, analyzed by CellChat. (**C**) Plots showing the expression of *Osm* and *Ifng* in young and aged TRL subsets. (**D**) Images of organoids derived from AT2s of young mice co-cultured with TRL subsets from young or aged mice. (**E** and **F**) Quantification of colony-forming efficiency (CFE) and organoid size. (**G**) Images of organoids derived from AT2s of young mice treated with recombinant OSM or IFNγ. (**H** and **I**) Quantification of colony-forming efficiency (CFE) and organoid size.

Furthermore, *Ifng* was preferentially expressed by CD4⁺ and CD8⁺ TRLs, while *Osm* was mainly expressed by DNTs (Fig. 5C). The upregulation of these two cytokines within TRL subsets of aged lungs was further validated by qPCR analysis (Fig. S4, A and B). Re-analysis of publicly available datasets (Table S6) (*29*) showed that *OSM* and *IFNG* were highly expressed in T cells of aged human lungs as well (Fig. S4C). Genes encoding the receptors for OSM and IFNγ were highly expressed by both mouse and human AT2s (Fig. S4, D and E). These data suggest that the T cell compartment engages in strong interactions with AT2s and that the upregulation of *Osm* and *Ifng* expression within T cells is conserved across mice and humans during aging.

To directly evaluate the suppressive capacity of each TRL subset, we co-cultured young AT2s with CD4^+^, CD8^+^, and DNT TRLs isolated from either young or aged mice using our organoid system (*9*). We found that all three TRL subsets from both young and aged lungs inhibited AT2 organoid formation, with this suppression being more pronounced in aged TRL subsets (Fig. 5, D-F). Furthermore, direct addition of recombinant OSM or IFNγ to AT2 organoids recapitulated the suppression of AT2 growth (Fig. 5, G-I). Together, these data indicate that all TRL subsets in aged lungs acquire an enhanced capacity to suppress AT2 growth relative to their counterparts in young mice, thereby positioning TRLs as key cellular mediators of age-associated emphysematous changes, at least in part through the secretion of OSM and IFNγ.

### IL-7R blockade limits TRLs and rescues emphysema in aged lungs

We have previously shown that lung fibroblasts create a niche that maintains TRL homeostasis, and that dysregulation of this niche promotes TRL expansion, thereby contributing to chronic lung disease (*9*). Here, we investigated how aging alters stromal-immune interactions in the lung by focusing on the crosstalk between the aged stromal niche and TRLs. To this end, we analyzed scRNA-seq data of stromal cells from young and aged lungs and categorized them into 5 distinct cell clusters (Fig. 6A). We next employed NicheNet, a tool for identifying and analyzing ligand-receptor interactions (*30*), to investigate intercellular communication between alveolar fibroblasts and TRLs, thereby assessing the capacity of the stromal niche to recruit and maintain TRLs within the alveoli. Notably, the top activated ligands were predominantly associated with T cell activation and cytokine secretion, including adrenomedullin (Adm) (*31*), bone morphogenetic protein 2 (Bmp2) (*32*) and haptoglobin (Hp) (*33*), as well as interleukin-15 (*Il15*) and interleukin-7 (*Il7*) (Fig. 6B), both of which are essential for TRL expansion and maintenance (*34*).

**Fig. 6.**
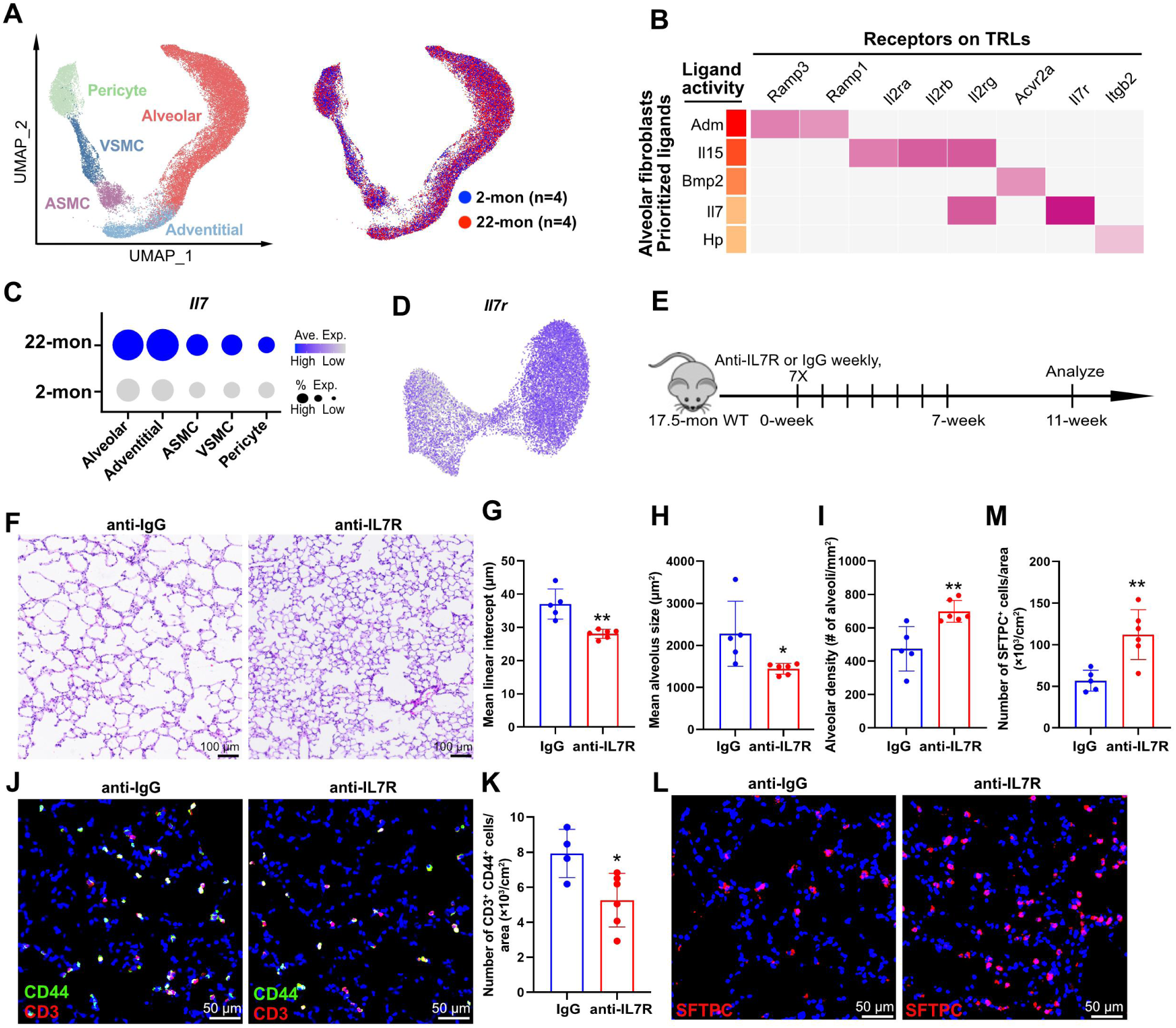
Limiting TRLs via IL-7R blockade restores alveolar integrity in aged lungs. (**A**) scRNA-seq of stromal cells from young and aged mouse lungs. (**B**) NicheNet interactome analysis of DEGs in TRLs (aged vs. young lungs) and alveolar fibroblasts. *Adm*, Adrenomedullin; *Il15*, Interleukin-15; *Bmp2*, Bone morphogenetic protein 2; *Il7*, Interleukin-7; *Hp*, Hephaestin. (**C**) Plots showing the expression of *Il7* in stromal subsets from young and aged lungs. (**D**) Plots showing the expression of *Il7r* in TRLs. (**E**) Schematic of aged mice treated with IgG isotype or anti-IL7R experiment. (**F**) H&E images of treated lungs. (**G** to **I**) Quantification of MLI, mean alveolus size, and alveolar density in treated lungs. (**J**) IF analysis of double-positive (CD3^+^/CD44^+^) T cells in treated lungs. (**K**) Quantification of double-positive (CD3^+^/CD44^+^) T cells in treated lungs. (**L**) IF analysis of SFTPC in treated lungs. (**M**) Quantification of AT2s in treated lungs. Each data point represents one mouse (**G** to **I**, **K**, **M**) of an individual experiment. Data are expressed as Mean ± SD. *p < 0.05; **p < 0.005; ****p < 0.0001.

Specifically, IL-15 has been shown to selectively promote the proliferation and survival of CD8^+^ TRLs, whereas IL-7 exerts a broader effect, supporting the maintenance of both CD4^+^ and CD8^+^ TRLs in tissues (*35–37*). Consistent with the NicheNet data, *Il7* displayed age-associated upregulation in stromal populations, including alveolar fibroblasts, and *Il7r* was broadly expressed across all TRL subsets (Fig. 6, C and D). We and others previously demonstrated that blockade of IL-7R signaling reduced the number of TRLs (*9, 38, 39*). Building on these findings, we administered weekly doses of anti-IL-7R antibody or isotype IgG to aged mice to assess whether IL-7R inhibition could mitigate age-associated emphysematous changes (Fig. 6E).

Histological evaluation of alveolar morphometry revealed that anti-IL-7R treatment attenuated alveolar space enlargement (Fig. 6, F-I), indicating a protective effect on alveolar integrity in aged lungs. Consistently, immunofluorescence analysis demonstrated that anti-IL-7R treatment decreased the number of TRLs (Fig. 6, J and K), while increasing the abundance of AT2s (Fig. 6, L and M). Collectively, our data suggest that the aged lung stromal niche promotes TRL expansion, leading to pathological suppression of AT2 function and the subsequent emphysema, whereas blocking the IL-7/IL-7R signaling axis reverses the emphysematous changes.

## DISCUSSION

Physiological lung aging is accompanied by progressive airspace enlargement and loss of alveolar surface area (*2, 40*), yet the causal immune-epithelial interactions that convert “inflammaging” into structural degeneration remain poorly defined. Here, we identify an age-associated expansion of TRLs as a previously underappreciated cellular driver of the emphysema phenotype in aging, extending prevailing models that have largely attributed this condition to age-related deterioration of the lung parenchymal supporting structure (*41*). Aging lungs exhibit a pronounced decline in the regenerative capacity of AT2s, concomitant with an expansion of TRLs exhibiting hallmarks of cellular senescence, including increased SASP (Fig. S5). Our data support a model in which these senescent TRLs remodel the alveolar niche to impair epithelial maintenance, thereby mechanistically linking immune aging to structural deterioration of the lung. Importantly, we further demonstrate that targeting of the IL-7/IL-7R axis limits TRL accumulation and partially preserves alveolar integrity in aged lungs, highlighting a potentially tractable immunomodulatory strategy to mitigate lung aging.

Our data further link TRL accumulation to immunosenescence. TRLs in aged lungs exhibit increased expression of canonical senescence markers, including p16, together with a SASP-like cytokine profile and activation of antigen-responsive signaling pathways. In parallel, we observe contraction of the T cell receptor repertoire and clonal expansion. Collectively, these features are consistent with chronic antigenic stimulation and sustained, tissue-embedded inflammatory signaling as defining components of immune aging (*42–44*). Elucidating the upstream drivers of clonal TRL expansion, including persistent infection, altered self-recognition, epithelial barrier dysfunction, or age-associated microbiome shifts, will be essential for identifying the initiating events that predispose the aged alveolar niche to degeneration.

Importantly, we provide in vivo evidence that aged lymphocytes are sufficient to promote alveolar destruction and AT2 loss. Adoptive transfer of T cells into lymphocyte-deficient recipients induced emphysematous changes, with aged donor cells producing more severe pathology. While this model cannot fully recapitulate the complexity of endogenous tissue residency, it supports a causal role for lymphocyte-intrinsic aging features in promoting alveolar decline. This finding is conceptually aligned with emerging studies showing that transplanting aged or senescent immune cells can impose tissue dysfunction across multiple organs in otherwise young hosts (*45–47*). Together with our in vitro data showing direct suppressive effects of TRLs on AT2 function, these in vivo findings reinforce a model in which immune cell aging actively drives deterioration of the alveolar epithelial compartment.

Finally, our work suggests a tractable therapeutic entry point at the level of niche maintenance rather than single cytokines. Consistent with prior observations that the aged lung environment promotes exaggerated accumulation of TRLs (*19*), our stromal-immune interactome analyses indicate that aging enhances T cell-supportive cues, including IL-7, while TRLs broadly express IL-7R. In line with this niche-centered framework, local IL-7R blockade attenuated TRL accumulation and ameliorated multiple morphometric features of age-associated emphysema, including restoration of AT2 abundance and alveolar density. This positions the IL-7/IL-7R axis as an upstream regulator of a chronic, TRL-driven cytokine milieu, including IFNγ and OSM, that constrains alveolar regeneration during aging.

Several limitations of this study warrant consideration. The analysis of human samples is constrained by cohort size, and while organoid systems enable mechanistic interrogation, they do not fully recapitulate the biophysical, spatial, and multicellular complexity of the aging alveolar niche. In addition, IL-7R signaling plays a central role in immune homeostasis, underscoring the need for future studies to delineate the balance between local therapeutic efficacy and potential costs to host defense in aged organisms. It will also be important to determine whether targeting downstream effectors such as IFNγ and OSM, either individually or in combination, can confer regenerative benefit while minimizing immunological risk. Collectively, our findings support a model in which age-associated alterations in the stromal niche sustain TRL accumulation and senescent-like activation, and these TRLs impose a cytokine-mediated constraint on AT2-dependent alveolar maintenance, thereby promoting the development of emphysema.

## MATERIALS AND METHODS

### Animal studies

INKBRITE (*p16^GFP/+^*) mice were generously provided by Dr. Tien Peng (*24*), and the *Sftpc^CreER^* line by Dr. Hal Chapman (*48*). *Rag1*^−/−^ and C57BL/6J mice were purchased from GemPharmatech and Shanghai Model Organisms, respectively. Both male and female animals were used in all experiments, as sex was not considered a biological variable.

### Human lung samples

Human lung tissues were obtained from distal regions of donor lungs that were deemed unsuitable for transplantation from brain-dead donors. Samples were stratified by age into young (42-45 years, n = 3) and aged (55-70 years, n = 3) groups.

### Animal treatment

To activate CreER, tamoxifen (Cat# T006000; TRC) was dissolved in corn oil and administered intraperitoneally at 200 mg/kg body weight per day for 3 days. CD3^+^ T cells were isolated from the spleens of young (2 months) or aged (19 months) C57BL/6J mice using Selleck Mouse CD3^+^ T Cell Isolation Kit (Cat# B90021, Selleck). CD3^+^ T cells from young or aged donors were adoptively transferred into 3-month-old *Rag1*^−/−^ mice by retro-orbital intravenous injection. To limit TRL expansion in vivo, anti-mouse IL-7R (Cat# BE0065, BioXCell) or IgG isotype control (Cat# BE0089, BioXCell) was administered intranasally at 100 μg/dose weekly per animal for 7 doses. At indicated time points, lungs were collected for flow cytometry or histological analysis.

### Histology and immunofluorescence

The right ventricle of the mice was perfused with 1-3 mL PBS and the lungs were inflated with 4% PFA in PBS, and then fixed in 4% PFA overnight at 4°C. After fixation, the lungs were washed with cold PBS 4 times for 30 minutes each at 4°C, and then dehydrated in a series of increasing ethanol concentration washes (30%, 50%, 70%, 95% and 100%) for a minimum of 2 hours per wash. The dehydrated lungs were incubated with xylene for 1 hour at RT and then with paraffin at 65°C twice for 90 minutes, and then embedded in paraffin. The lungs were sectioned at 8 µm on a microtome. For frozen OCT-embedding, lungs were inflated and fixed with 4% PFA for 2 hours at 4°C, washed with PBS 4 times for 30 minutes each at 4°C, and embedded in OCT after 30% sucrose incubation.

For immunofluorescent staining, paraffin sections were incubated in xylene twice for 10 minutes, then rehydrated in ethanol washes (100% twice, 95%, 70%, 50% ethanol) for 5 minutes each. OCT embedded slides were fixed in 4% PFA at RT for 10 minutes, then washed three times with PBS. For both paraffin and OCT embedded slides, antigen retrieval (Cat# BRR2004CLX; Biocare Medical) was performed for 30 minutes at 95°C. Slides were washed with 0.1% Tween 20 in PBS (PBST) 3 times for 5 minutes. Slides were then incubated in blocking buffer (3% donkey serum in PBST) for at least 1 hour, then incubated overnight in primary antibodies in 1% donkey serum in PBST at 4°C. The following primary antibodies were used for mouse tissue, with the exception of rabbit anti-SPC (Cat# AB3786; Millipore Sigma; used 1:2000), which was applied to both mouse and human tissues: rabbit anti-CD3 (Cat# ab16669; Abcam; used 1:400), rat anti-CD44 (Cat# 103001; BioLegend; used 1:200), rat anti-CD45 (PE; Cat# 553081; BD; used 1:200), rat anti-CD45 (APC; Cat# 559864; BD; used 1:200). The following secondary antibodies were used at 1:500: donkey anti-rabbit Alexa Fluor 488 (Cat# A-21206; Thermo Fisher), donkey anti-rabbit Alexa Fluor 555 (Cat# A-31572; Thermo Fisher), donkey anti-rabbit Alexa Fluor 647 (Cat# A-31573; Thermo Fisher), donkey anti-rat Alexa Fluor 647 (Cat# 712-605-153; Jackson ImmunoResearch), and donkey anti-rat Alexa Fluor 555 (Cat# A-78945; Life Technologies).

DAPI (0.2 µg/mL) (Cat# XW287189031; Sinopharm) was added for 5 minutes, and after washing three times with PBS for 5 minutes each, slides were mounted using FluorSave™ reagent (Merck) to maintain fluorescence.

### Lung digestion and cell isolation

For fluorescence activated cell sorting (FACS) analysis of immune cells, a digestion cocktail of Collagenase Type I (Cat# 17100017; Thermo Fisher; used 700 U/mL) and Dnase I (Cat# DN25; Sigma; used 50 U/mL) was used to dissociate the lung. The lung was further diced with a razor blade and incubated in digestion cocktail for 25 minutes at 37°C with continuous shaking. The mixture was then washed with sorting buffer (2% FBS (Cat# 10091148; Thermo Fisher) and 1% Antibiotic-Antimycotic (Cat# 15240062; Thermo Fisher) in DMEM (Cat# 31053028; Thermo Fisher)). The mixture was passed through a 70 µm cell strainer and resuspended in RBC lysis buffer (Cat# R7757; Sigma), then passed through a 40 µm cell strainer. Cell suspensions were incubated with the appropriate conjugated antibodies in sorting buffer for 30 minutes at 4°C and washed with sorting buffer. FACS was performed on a CytoFLEX SRT using CytoExpert SRT software. Doublets were excluded based on forward and side scatter, and dead cells were excluded by DAPI staining. The following antibodies were used for staining: CD45 (APC; Cat# 559864; BD; used 1:200), CD45 (PerCP/Cyanine5.5; Cat# 103132; BioLegend; used 1:200), CD4 (PE/Cyanine7; Cat# 100422; BioLegend; used 1:200), CD8a (BV605; Cat# 100743; BioLegend; used 1:200), CD3e (BV421; Cat# 100228; BioLegend; used 1:200), NK1.1 (BV785; Cat# 156539; BioLegend; used 1:200), CD19 (APC/Cyanine7; Cat# 115530; BioLegend; used 1:200), CD3e (FITC; Cat# 100306; BioLegend; used 1:200), Ly-6G (APC/Cyanine7; Cat# 127624; BioLegend; used 1:200), CD11b (BV510; Cat# 101263; BioLegend; used 1:200), CD11c (BV785; Cat# 117335; BioLegend; used 1:200), MHCII (Pacific Blue; Cat# 107620; BioLegend; used 1:200), B220 (APC-A700; Cat# 103232; BioLegend; used 1:200), Ly-6C (BV570; Cat# 128029; BioLegend; used 1:200), MerTK (PE/Cyanine7; Cat# 151522; BioLegend; used 1:200), CD64 (BV650; Cat# 139337; BioLegend; used 1:200), SiglecF (BV711; Cat# 155537; BioLegend; used 1:200). Analysis was performed using FlowJo v10 software. For distinguishing the resident and circulating lymphocytes, 5 min before euthanizing each mouse, 3 mg of anti-mouse fluorophore-conjugated CD45 antibody diluted in 100 μL of 1× DPBS was intravenously injected through retro-orbital injections.

For scRNA-seq encompassing diverse cell lineages, the whole mouse lung was dissected and tracheally perfused with a digestion cocktail of Dispase II (Cat# 17105041; Thermo Fisher; used 15 U/mL), Collagenase Type I (Cat# 17100017; Thermo Fisher; used 500 U/mL) and Dnase I (Cat# DN25; Sigma; used 50 U/mL), and removed from the chest.

### Single cell capture and sequencing

For scRNA-seq, all live lung non-immune cells and ivCD45-negative immune cells were sorted from young or aged C57BL/6J mice. Lung non-immune cells were isolated based on forward and side scatter, DAPI (Cat# XW287189031; Sinopharm; 0.2 µg/mL), CD31 (AF488; Cat# 102414; BioLegend; used 1:200) and CD45 (PE; Cat# 13-0451-85; Invitrogen; used 1:200) exclusion.

IvCD45-negative immune cells were isolated based on DAPI, CD31 exclusion, and ivCD45 (APC; Cat# 559864; BD; used 1:200) negative selection. The sorted cells were then loaded onto a single lane per sample in the Chromium^TM^ Controller to produce gel bead-in emulsions (GEMs). GEMs underwent reverse transcription for RNA barcoding and cDNA amplification, with the library prepped using the Chromium Next GEM Single Cell 5’ Kit v2. Each sample was sequenced on a single lane of the Illumina NovaSeq 6000.

### Organoid assay

For mouse AT2 only organoid assay, AT2s were isolated from C57BL/6J mice by gating based on CD326 (BV786; Cat# 740958; BD; used 1:200) and CD104 (PerCP/Cyanine5.5; Cat# 123614; BioLegend; used 1:200). tdT^+^ AT2s were sorted from tamoxifen-injected SftpccreERT2/+:R26RtdTomato mouse lungs. Then AT2s were resuspended in a feeder-free medium modified from the previously reported study (*49*). Briefly, advanced DMEM/F12 medium was supplemented with 10 μM SB431542 (Cat#Ab120163; Abcam), 3 μM CHIR (Cat#4423-10; Tocris), 1 μM BIRB796 (Cat#598910; Tocris), 50 ng/mL mouse EGF (Cat#HY-P7194; MCE), 10 ng/mL mouse FGF10 (Cat#10573-HNAE-20; SinoBiological), 10 ng/mL mouse IL-1β (Cat#579402; BioLegend), 5 μg/mL Heparin (Cat#9041-08-1; Sigma), 1× B-27 supplement (Cat#17504001; Gibco), 1× Antibiotic-Antimycotic (Cat#15240062; Gibco), 15 mM HEPES (Cat#931621681; Yuanyebio), 1× GlutaMAX (Cat#35050061; Gibco), 1.25 mM N-Acetyl-L-cysteine (Cat#A9165; Sigma), and 10% FBS (Cat#A5669701; Gibco). The resuspended AT2s were mixed with an equal volume of Matrigel (Cat#354230; Corning). 50 μL of the mixture (1× 10^4^ AT2s) was added to each well of 24-well plate, and 500 μL medium was added to each well after Matrigel solidification. For mouse AT2-TRL coculture, 3×10^3^ TRLs were added into the well of 24-well plate with 10 μL mouse T-Activator CD3/CD28 beads (Cat#11456D; Thermo Fisher), and the medium was supplemented with 100 U/mL human IL2 (Cat#402-ML-020; R&D). Where applicable, 1 ng/mL recombinant mouse IFNγ (Cat#BE0055-1MG; BioXcell) and 20 ng/mL recombinant mouse OSM (Cat#50112-M08H; SinoBiological) were added to the medium. 10 μM Y-27632 (Cat#S1049; Selleck) was added to the medium for the initial 3 days. The medium was changed every 3-4 days. The mouse organoids were analyzed at day 12-14.

### Quantitative PCR

Total RNA was obtained from cultured cells using the FastPure Cell/Tissue Total RNA Isolation Kit V2 (Cat# RC112; Vazyme), following the manufacturer’s protocol. cDNA was synthesized from total RNA using the HiScript III RT SuperMix (Cat# R323; Vazyme). Quantitative PCR (qPCR) was performed using the SYBR Green system (Cat# Q711; Vazyme). Primers are listed in Table S7. Relative gene expression levels after qPCR were defined using the ΔΔCt method and normalizing to *GAPDH*.

### Immunofluorescence image quantification

Sections were imaged on the Olympus FV3000 or Zeiss LSM 980 Confocal Microscope for quantification. At least three samples per genotype/condition were used, and at least 5 randomly selected sections were chosen for each sample. Cell counts for stained cells were performed in Fiji using the “Cell Counter” plug-in. For quantification by area, the image was first converted to 8-bit and the “Measure” function was used. For all analyses, performer was blinded to the specimen genotype/condition during data collection and analysis. Results were averaged for each specimen and standard deviations were calculated per genotype/condition. Images of CD3^+^/CD44^+^ (mouse lungs) co-staining were processed in CellProfiler (Broad Institute). Primary objects were generated using the Identify Primary Objects module for each fluorescent channel. Size and mean intensity filters were applied to remove debris. Double-positive cells were identified by overlapping primary objects with the Relate Objects module. Double-positive cells were used to generate a mask that was applied to both channels of the original image file.

### 3D image analysis and quantification

3D image analysis was performed as previously described (*50, 51*). Briefly, z-stack images were rendered in 3D dimensions and quantitatively analyzed using Bitplane Imaris v9.5 software package (Andor Technology PLC, Belfast, N. Ireland). Individual ivCD45^−^/CD3^+^ lymphocytes were annotated using the Imaris spots function based on the colocalization analysis.

### MLI, airspace size, and alveolar density analysis

For alveolar morphometric analysis, lungs were processed according to the above protocol for paraffin-embedded samples, with the exception of inflation with 4% PFA at a constant pressure of 25 cm H2O. The paraffin-embedded lung sections were stained by hematoxylin and eosin (H&E) for analyzing alveolar morphology metric. At least 5 randomly selected sections from each genotype were selected for analysis. The mean linear intercept (MLI) was calculated as the linear sum of the lengths of all lines randomly drawn across the images, divided by the number of intersections between alveolar walls. A minimum of 1000 intercepts from 50 lines drawn across the lung in a randomized fashion were obtained for each lung, and the analysis was carried out on Fiji with the “Cell Counter” plug-in. The airspace size was measured with Fiji using the Analyze Particle tool. Images were first converted to 8-bit and inverted, and the “Analyze Particle” function was used with a set minimum of 50 µm^2^ and maximum of infinity to identify and quantify alveoli in the image. The average airspace size for each lung was quantified by dividing the total airspace by the number of alveoli. At least 3,500 alveoli were measured for each lung.

The alveolar density was the reciprocal of the airspace size. The airspace of airway and pulmonary vessels was excluded.

### Single cell RNA-seq analysis

For the animal data, FASTQ files were run through CellRanger version 6.1.0 software with default settings for de-multiplexing, aligning reads with STAR software to mm10, and counting unique molecular identifiers (UMIs). Seurat package v5 in RStudio was used for downstream analysis (*52*). Low-quality cells were filtered (expressing fewer than 200 genes, more than 10% mitochondrial reads, or more than 6,000 unique gene counts). Principal component analysis was performed on log-normalized and scaled data using 2,000 variable genes. The top 30 principal components were used for clustering and visualized using the UMAP algorithm in the Seurat package. The lists of DEGs were identified with a Model-based Analysis of Single-cell

Transcriptomics (MAST) test. Pathway and Upstream Regulator analyses of gene lists containing significantly differentially expressed genes were done with Ingenuity Pathway Analysis (Qiagen). The representation factor was calculated to represent the number of overlapping genes divided by the number of expected overlapping genes drawn from two independent groups, as calculated on nemates.org with a base value of 30,000 genes in the genome. For interactome analysis we used the NicheNet and CellChat packages.

### Statistical analysis

All statistical analyses were performed in GraphPad Prism 8.0.2 using unpaired one-tailed Student’s t-tests to determine the P-value. Specific *P* values were labeled in the plots or figure legends, where significant values were *P* < 0.05. All data in graphs were presented as mean ± SD. No data were excluded from the analyses presented in this study. Three or more replicates were used for each experiment, and the specific n values were provided in the figure legends.

## Acknowledgments

We thank Dr. Tien Peng for sharing INKBRITE (*p16^GFP/+^*) mice and Dr. Hal Chapman for the *Sftpc^CreER^* mice.

## Author contributions

Conceptualization: Y.S., X.Y., Z.D., C.W.

Investigation: Y.S., X.Y., Z.D., C.W.

Data curation: Y.S., Z.R., Y.G, C.W.

Validation: Y.S., X.Y., X.Z., S.C., Y.H.

Formal analysis: Y.S., X.Y., T.Y., J.L.

Visualization: Y.S., X.Y., Y.G, Z.R.

Resources: F. G., D.C., J.C., Z.D., C.W.

Project administration: F. G., D.C., J.C., Z.D., C.W.

Supervision: F. G., D.C., J.C., Z.D., C.W.

Funding acquisition: C.W. Writing—original draft: Y.S., Z.D., C.W.

Writing—review & editing: Y.S., F. G., D.C., J.C., Z.D., C.W.

## Data and materials availability

All data are available in the main text or the supplementary materials. The sequencing data reported in this study are deposited in NCBI Gene Expression Omnibus (GEO) under the accession number: GSE309583. The information of previously published scRNA-seq data re-analysed here is available in the Supplementary Information. This paper does not report any original code.

