## Supplemental material for "Dysregulated tissue-resident lymphocytes drive senile emphysema by impairing alveolar regeneration"

Yanjing Su *et al.*

*Corresponding author. Email:
 (F.G.)
 (D.C.)
 (J.C.)
 (Z.D.)
 (C.W.)

**This PDF file includes:**

Figs. S1 to S5
Tables S1 to S7

**Fig. S1.**

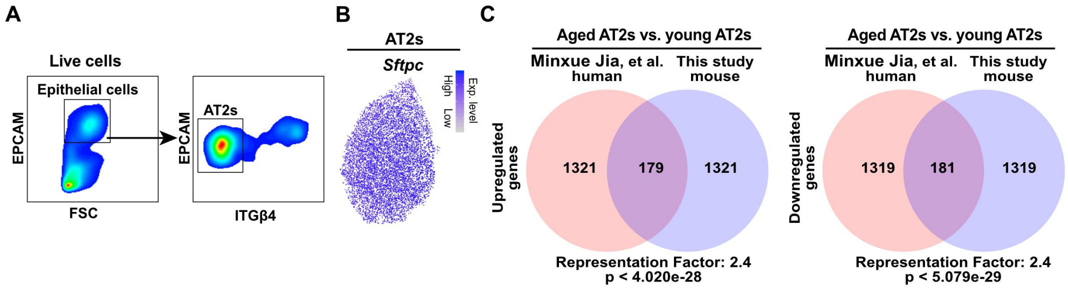


**Fig. S1. Conserved age-associated transcriptional remodeling of AT2s in humans and mice.** (**A**) FACS strategy to isolate AT2s from wild-type mouse lungs. (**B**) Plots showing the expression of AT2 markers, including *Sftpc*. (**C**) Comparison of the top 1000 DEGs in human (aged vs. young AT2s) and mouse (aged vs. young AT2s), with statistical comparison for degree of gene overlap of two independent gene sets denoted by representation factor (see Methods).

Fig. S2.

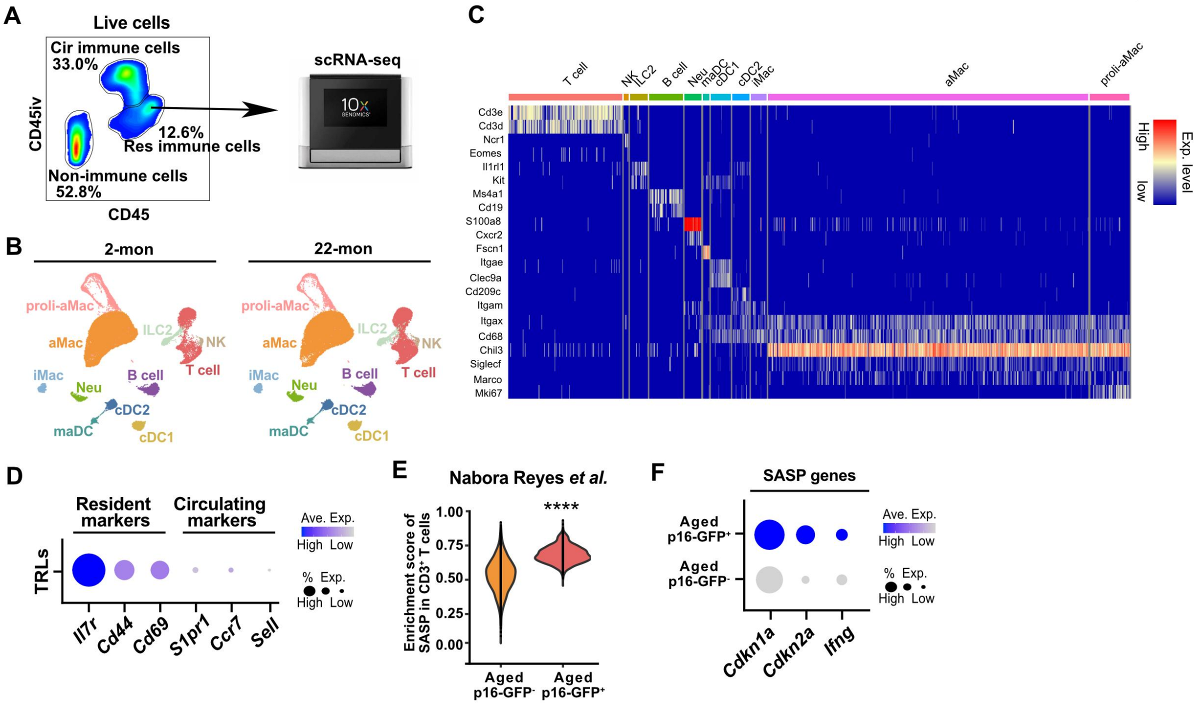


**Fig. S2. Expansion TRLs with SASP signature in aged lungs.**

(**A**) FACS strategy to isolate resident immune cells for scRNA-seq. (**B**) UMAP plots showing cell clusters in young (2-month-old, n = 7) and aged (22-month-old, n = 7) lungs. (**C**) Heatmap showing marker genes for each resident immune subset. (**D**) Plots showing the expression of resident markers (*Il7r*, *Cd44*, and *Cd69*) and circulating markers (*S1pr1*, *Ccr7* and *Sell*) in TRLs. (**E**) Plots showing the enrichment score of SASP in young and aged $CD3^+$ T cells isolated from INKBRITE ($p16^{GFP/+}$). (**F**) Plots showing the expression of SASP genes in young and aged $CD3^+$ T cells isolated from INKBRITE ($p16^{GFP/+}$).

**Fig. S3.**

A

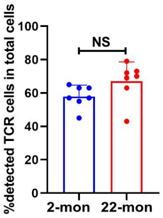


**Fig. S3. High frequency of paired TCRαβ TRLs in young and aged lungs.**

(**A**) Percentage of TRLs assigned paired TCRα and TCRβ chains in young and aged lungs. Each data point represents one mouse of an individual experiment. Data are expressed as Mean ± SD. $*p < 0.05$; $**p < 0.005$; $****p < 0.0001$.

### Fig. S4.

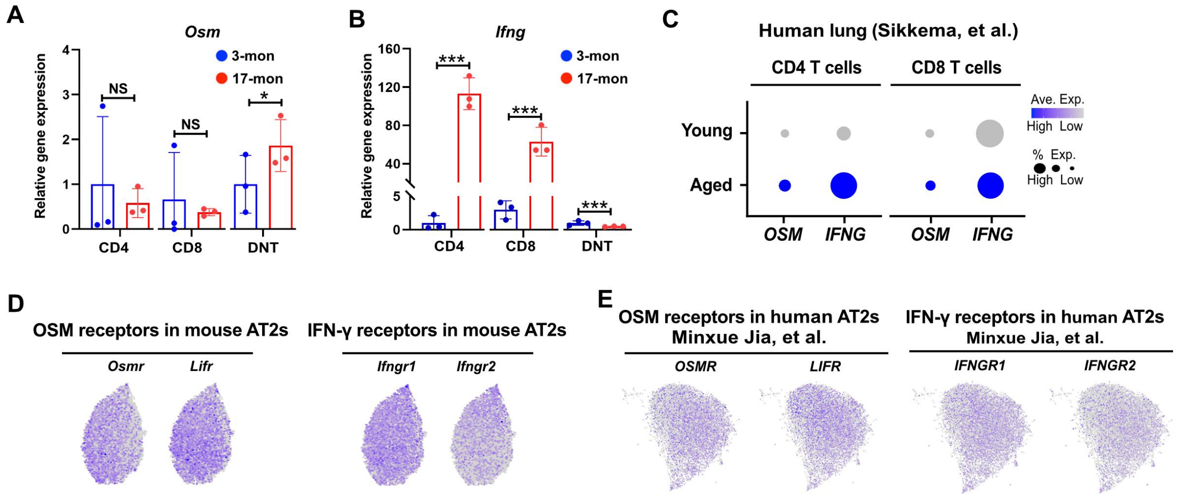


**Fig. S4. Expression of OSM, IFNγ, and their receptors in humans and mice.**

(**A** and **B**) qPCR analysis of *Osm* and *Ifng* expression in TRL subsets isolated from AT2-TRL organoid co-cultures. (**C**) Plots showing the expression of *OSM* and *IFNG* in $CD4^+$ or $CD8^+$ T cells from young and aged human lungs. (**D**) Plots showing the expression of *Osm* and *Ifng* receptors in AT2s from young and aged mouse lungs. (**E**) Plots showing the expression of *OSM* and *IFNG* receptors in AT2s from young and aged human lungs.

**Fig. S5.**

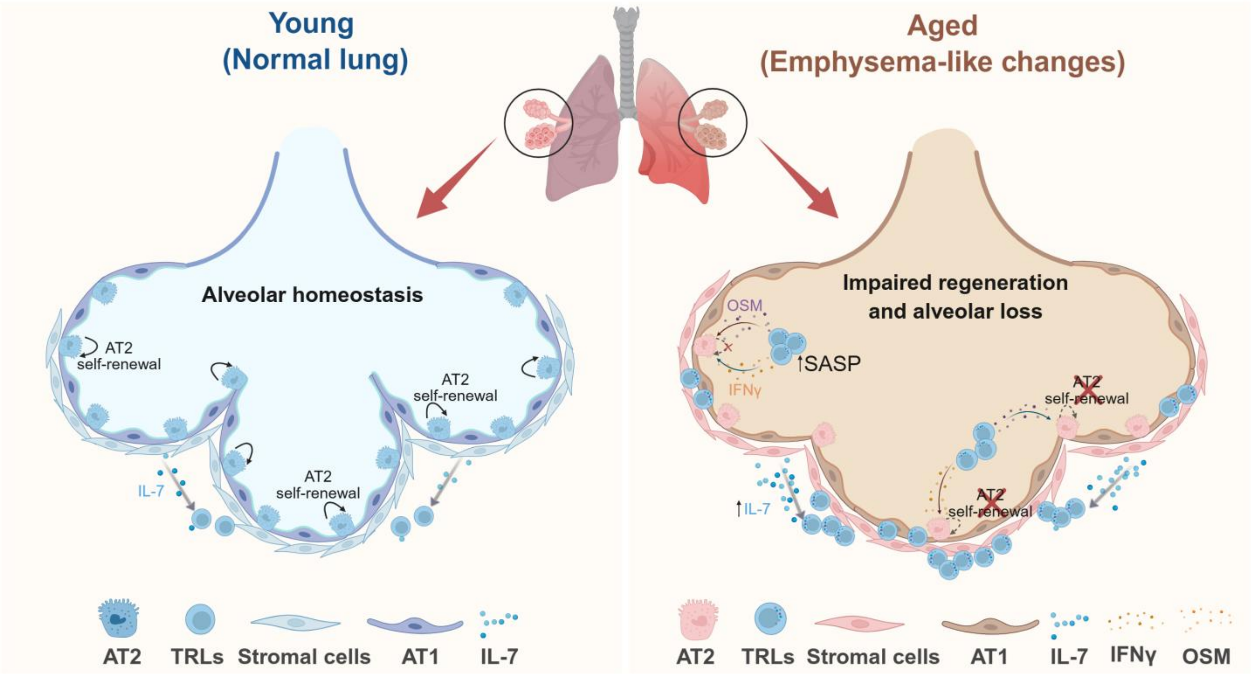


**Fig. S5. Model of tissue-resident T lymphocytes (TRLs) driving age-associated emphysema.**

**Table S1. Human lung information (Histology and immunofluorescence)**

**Table S2. Differentially expressed genes in aged vs young mouse AT2**

**Table S3. Public scRNA-seq datasets for comparing aged and young human AT2s**

**Table S4. Differentially expressed genes in aged vs young human AT2**

**Table S5. Differentially expressed genes in aged vs young mouse TRLs**

**Table S6. Public scRNA-seq datasets for comparing aged and young human T cells**

**Table S7. qPCR primers**
